# Survival cost to relocation does not reduce population self-sustainability in an amphibian

**DOI:** 10.1101/446278

**Authors:** Hugo Cayuela, Lilly Gillet, Arnaud Laudelout, Aurélien Besnard, Eric Bonnaire, Pauline Levionnois, Erin Muths, Marc Dufrêne, Thierry Kinet

## Abstract

1. Relocations are increasingly popular among wildlife managers despite sharp debate and low rate of relocation success in vertebrates. In this context, understanding the influence of extrinsic (e.g., relocation design, habitat characteristics) and intrinsic factors (e.g., age and sex) on demographic parameters such as survival that regulate the dynamics of relocated populations is critical to improve relocation protocols and better predict relocation success.
2. We investigated survival in naturally established and relocated populations of yellow-bellied toads (*Bombina variegata*), an amphibian that was nearly extinct in Belgium by the late 1990s. We quantified survival at three ontogenetic stages (juvenile, subadult, and adult) in the relocated population, the source population, and a control population. In the relocated population, we quantified survival in captive bred individuals and their locally born descendants.
3. We showed that survival at juvenile and subadult stages was relatively similar in all populations. In contrast, relocated adult survival was lower than adult survival in the source and control populations. Despite this, offspring of relocated animals (the next generation, regardless of life stage) survived at similar rates to offspring in the source and control populations. Our simulations revealed that the relocated population was self-sustaining under different scenarios and that the fate (e.g., stability or finite rate of increase) of the simulated populations was highly dependent on the fecundity of relocated adults and their offspring.
4. *Policy implications*. Our results indicate that survival in relocated individuals is lower than in non-relocated individuals but that this cost (= reduced survival) disappears in the second generation. A finer understanding of how relocation affects demographic processes is an important step in improving relocation success of amphibians and other animals.

## Introduction

Evaluating the efficiency of wildlife management methods to mitigate population declines and reduce extinction risk are central goals for conservation biologists. Over the three last decades, methods termed relocation (sensu Fischer & Lindenmayer 2000, including the terms reintroduction, translocation, and supplementation), have become increasingly popular among wildlife managers (Seddon et al. 2007, 2012). These methods involve moving individuals or groups of individuals from free-ranging or captive populations to new or historical habitats, generally in response to threat (e.g., habitat loss, exotic species, disease, Fischer & Lindenmayer 2000, Seddon et al. 2007, Armstrong & Seddon 2008). The efficiency of relocation programs has been debated (Griffith et al. 1989, Hodder & Bullock 1997, Seddon et al. 2012), but in general a relocation is considered successful when it results in a self-sustaining (i.e., viable, with population growth ≥ 1) population (Seddon et al. 2007, 2012; Robert et al. 2015). In vertebrates, relocation success rate is often relatively low and depends on multiple factors ranging from conditions during captive breeding (e.g., diet, habitat, conspecific interactions; Jule et al. 2008; Whiteside et al. 2015) to relocation protocols (e.g., number of individuals released and the frequency of release, age or ontogenic stage at release; Griffith et al. 1989, Fischer & Lindenmayer 2000; Germano & Bishop 2009), to environmental characteristics (e.g., habitat quality, potential anthropogenic threats; Griffith et al. 1989; Parlato & Armstrong 2013) and biotic interactions (e.g. predation, competition: Fischer & Lindenmayer 2000; Parlato & Armstrong 2013) at the release site. Understanding the underlying mechanisms of why a particular translocation is successful is paramount to increasing the efficacy of relocation as a conservation tool (Converse et al. 2013).

One mechanism may be how these factors affect demographic parameters (e.g., survival). Understanding how demographics may change in relocated populations is understudied, yet is critical to identifying how management strategies can be modified to increase and better predict the success of relocation. For example, survival immediately following the release of animals into new habitat affects the outcome of a relocation (i.e., establishment phase, Armstrong & Seddon 2008, Armstrong et al. 2017). Reduced survival during the establishment phase occurs in a range of vertebrates (birds: Armstrong et al. 2017; mammals: Jule et al. 2008; reptiles: Roe et al. 2010, fishes: Jonsson et al. 2003), but the magnitude of this decrease varies among taxa (Jule et al. 2008, Armstrong et al. 2017) and is influenced by multiple factors. Physiological (chronic or acute) stress response, typically associated with animal capture, transport, and relocation can result in a reduced survival after relocation (Dickens et al. 2010, Tarszisz et al. 2014). Predation, interspecific competition, and mal-adaptation to novel environmental conditions can also negatively affect post relocation survival (Banks et al. 2002, Roe et al. 2010, Lovari et al. 2014). In contrast, some species, or individuals within a species, may have the capacity to respond to the novel environment and change their physiology, behaviour, synaptic connections, immune repertoire, or other characteristics in an adaptive direction (West-Eberhard 1989, Ghalambor et al. 2007, Fusco & Minelli 2010). Environmentally-induced changes in individual phenotype, called “acclimatization” or “acclimation”, can increase post-relocation survival (Armstrong et al. 2017). The ability of individuals to accommodate novel environmental conditions may also depend on individual factors such as sex and age (Teixeira et al. 2007). The effect of sex has been investigated (Pyne et al. 2010, Gouar et al. 2008, Armstrong et al. 2017), but reduced survival during the establishment phase and related consequences for relocation success remain poorly understood (but see Gouar et al. 2008, Aaltonen et al. 2009).

Additionally, there is a taxonomic bias in reports of post-relocation survival towards birds and mammals (Seddon et al. 2005, Bajomi et al. 2010). Notably, the study of the demographic processes related to relocation success has been neglected in amphibians (Griffiths & Pavajeau 2008, Germano & Bishop 2009; but see Bodinof et al. 2010, Muths et al. 2014, Howell et al. 2016). Not only are amphibians the most endangered vertebrate in the world (IUCN 2018), but relocation success in amphibians is substantially lower than in other vertebrates (Dodd & Seigel 1991, Griffiths et al. 1989).

Relocation projects have been undertaken in a broad range of amphibian taxa (with varying degrees of success, e.g. Muths et al. 2001, Seigel & Dodd 2002, Griffiths & Pavajeau 2008, Germano & Bishop 2009, Harding et al. 2016), and relocations, combined with captive breeding programs, have been proposed as promising management tools for some pond-breeding amphibians (Marsh & Trenham 2001, Griffiths & Pavajeau 2008). In fact, the number of captive breeding and release programs for amphibians is increasing globally (Germano & Bishop 2009, Harding et al. 2016) suggesting that interest is high in these methods, despite overall low success rates and some notable gaps in information. For example, few studies have investigated post-relocation demographic processes (but see Bodinof et al. 2010, Muths et al. 2014, Howell et al. 2016) such as the presence and consequences of variation in adult survival during the establishment phase or patterns in age-dependent survival in relocated populations.

To address this knowledge gap, we quantified survival rates in juvenile, subadult and adult yellow-bellied toads (*Bombina variegata*) in a relocated population, the source population (from which adults involved in the captive breeding program were collected), and a control population (demographically and genetically independent) in Belgium where this amphibian is nearly extinct (de Wavrin 2007). This enabled us to determine if age-dependent survival occurred and if there were differences in patterns of age-dependent survival among populations (e.g., differences in adult survival post-relocation). We used multievent CR models to estimate parameters and examine differences among populations. We hypothesized that (1) survival increases with age (i.e., age-dependent survival); (2) survival is different in relocated adults compared to adults in the source and control populations; and (3) survival is different between relocated individuals and their offspring (next generation). We also assessed the long-term viability of the relocated population by simulating population trajectories under several different scenarios focused on fecundity (i.e. the number of juvenile produced per female per year). We discuss our results in the context of the *B. variegata* captive breeding and relocation program and in the broader context of amphibian relocation efforts.

## Materials and methods

### Study area

Our study was conducted on three populations of *B. variegata* in Belgium and north-east France (**Fig.1**). The source population has been breeding in a set of artificial ponds in an old quarry since 1985. Individuals involved in the captive breeding were collected in this population and placed in ex-situ terraria (see breeding design, below).

**Fig.1.**
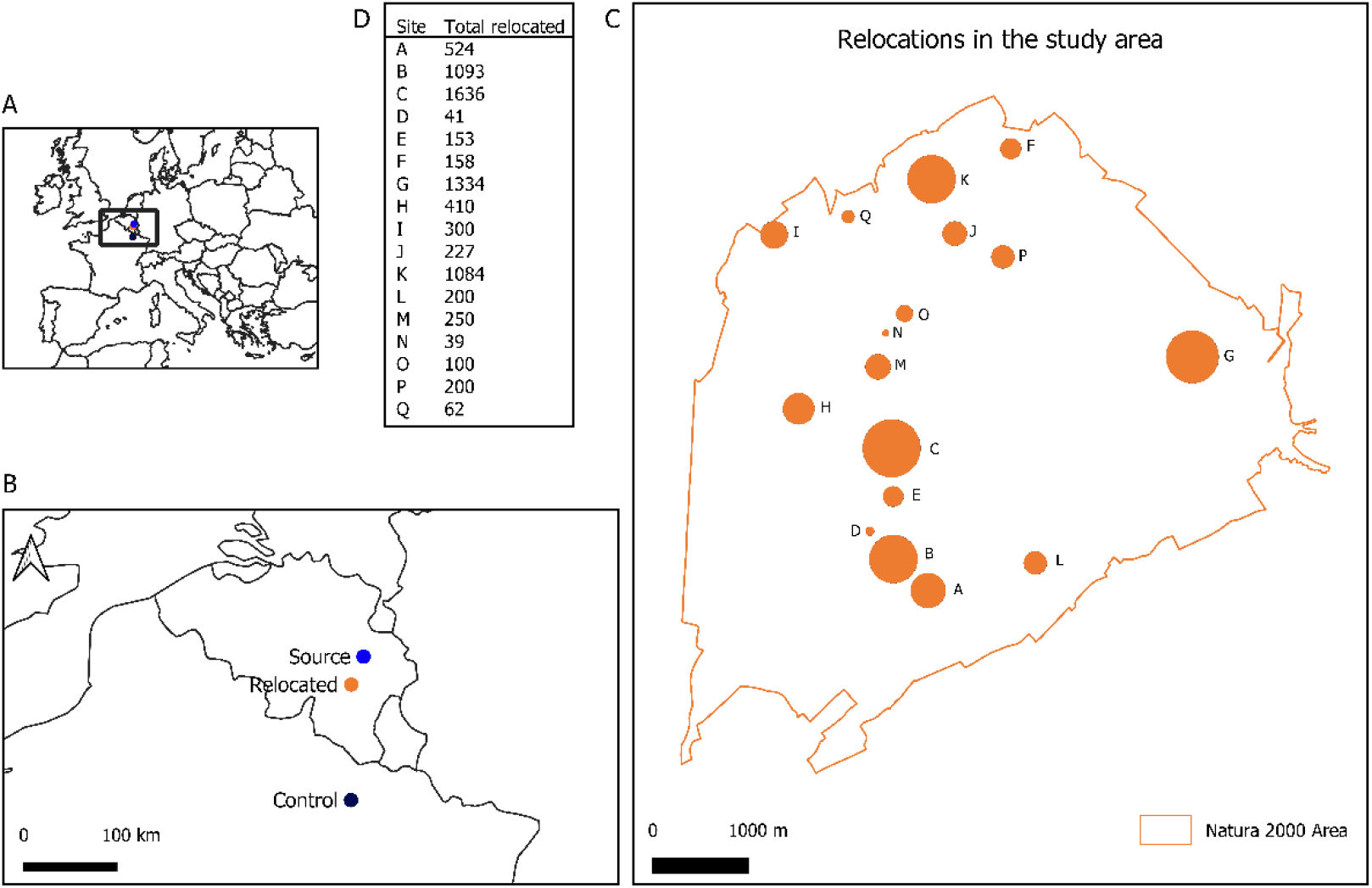
Map of the study area. Maps A and B show the localization of three studied populations (control, source and relocated). Map C shows the relocation sites (i.e. groups of breeding waterbodies) in the study area. Table B provides the total number of individuals relocated each relocation site.

Individuals from the captive breeding were relocated to a site at Marche-en-Famenne, Belgium (relocated population) which is an active military training field that is part of the Natura2mil Life Program. Unauthorized access to the site strictly forbidden, which avoids threats contributing to decline in *B. variegata* (i.e., habitat loss caused by agriculture and urban development Cayuela et al. 2015b, 2018b). The relocation site is a mosaic of grassland and woodland areas (Denoël et al. 2018). The relocated population breeds in 20 distinct groups of ruts separated by 692 to 4044 m (mean = 2336 m; **Fig. 1**). Dispersal between the groups of ruts is very low (only one individual of 635 captured have moved between 2 distinct sites over the study). Human activities at the site are limited to military vehicle traffic from late April to September, which includes the breeding season for *B. variegata.* Vehicle traffic creates breeding waterbodies and maintains the temporary nature of these ponds where *B. variegata* reproduces.

The control population is located in a harvested woodland in north-east France, close to Verdun (Cayuela et al. 2018a). This population reproduces in set of 189 sites (i.e., groups of ruts) and dispersal is high between the sites (Cayuela et al. 2016a).

### Captive breeding and relocation design

Thirty individuals were captured in the source population and placed in several ex-situ terraria (**Fig. 2**) where natural conditions were mimicked (rise of the water level) to trigger 3 or 4 breeding periods each year. Stocking at the relocation site lasted six years (2008-2014). At each breeding season, the eggs laid in the enclosure were collected and placed in outdoor tanks (from 40×35×20 to 400×120×40 cm). Tadpoles were fed *ad libitum* with boiled lettuce. When tadpoles reached the Gosner stages 36-45 (with the four legs developed; Gosner 1960), they were captured and released in the 20 groups of tuts at Marche-en-Famenne (**Fig. 2**). Each group of ruts was stocked from one year to four years (Appendix 1). We released 7804 tadpoles (Gosner stages 36-45) between 2008 and 2014 (**Fig. 2**). A few individuals (N = 18) were released at subadult (two years) and adult (three years or more) stages between 2009 and 2014 (**Fig. 2**).

**Fig.2.**
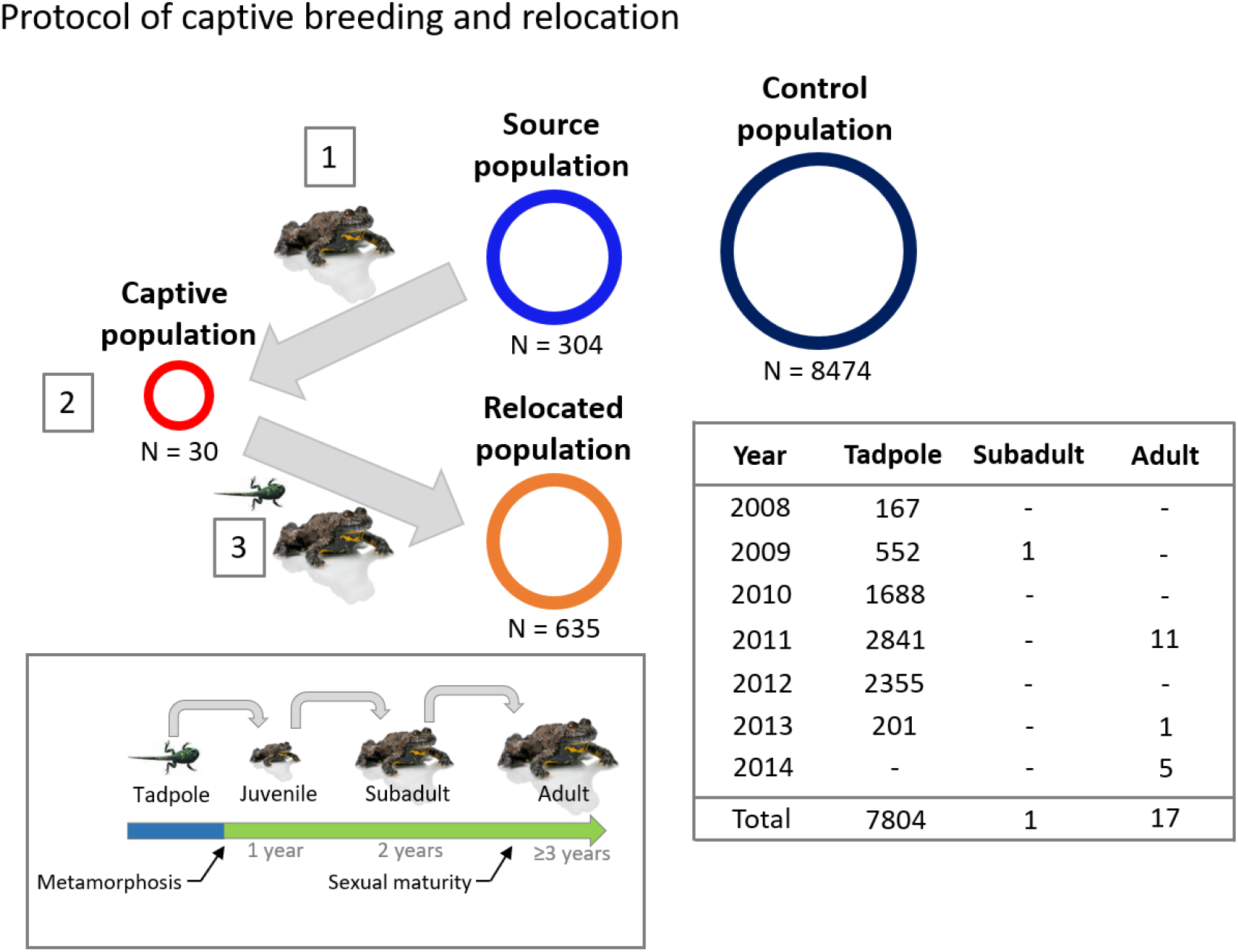
Protocol of captive breeding and relocation in Bombina variegata. This species has a complex life cycle: the reproduction takes place in waterbodies in which egg clutches are laid. After hatching, the tadpoles develop until metamorphosis (61 days at 27°C., 107 days at 15°C.; Morand et al. 1997). From metamorphosis to the first overwintering, individuals are terrestrial. After the first overwintering, individuals stare juveniles (1 year). If they survive, they reach the subadult stage after the second overwintering. After the third overwintering, individuals are sexually mature (Cayuela et al. 2017b). Juvenile and adult toads can be individually recognized and surveyed using capture-recapture methods (Cayuela et al. 2016b, 2016c). Before individuals reach 1 year, the aposematic colors used to recognize toads are not always sufficiently fixed so that individual recognition is not certain. Three populations of Bombina variegata were considered in our study: (1) a control population in eastern France, with a large size (8474 identified over 5-year study period), (2) the source population (304 individuals identified), the last natural population of B. variegata in Belgium, and (3) the relocated populations (635 individuals identified). For the captive breeding and the relocation, we had the following procedure: step 1, 30 individuals were collected from the source population and placed in ex-situ enclosures (i.e. captive population); step 2, captive individuals reproduce, eggs laid were collected and placed in tanks until tadpoles reached the Gosner stages 36-45. Individuals were released in the 20 sites (i.e. groups of ruts) inhabited by the relocated population at different stages (tadpoles at Gosner stages 36-45, subadult and adult). The stocking period lasted 6 years (2008-2014). The number of individuals released each year is shown in the table.

### Capture-recapture surveys

In the three populations, individuals were surveyed using capture-mark-recapture methods from 2008 through 2017. The relocated population was surveyed over the whole period (i.e., nine years), while control and source populations were monitored during five (2012-2016) and six years (2011-2016) respectively. The number of capture sessions varied among populations and years (**Supplementary material S1**). At each session, all suitable ruts were sampled exhaustively until no new individuals were detected. During sampling sessions, toads were caught by hand or net and were held boxes until photographed and released in their waterbody. Toads were assigned to one of three age classes (**Fig. 2**): Juveniles captured after their first overwintering, subadults (two overwinterings) and sexually mature adults (three or more overwinterings). We identified each individual by the unique pattern of black and yellow mottles on its belly and recorded these in photographs (**Supplementary material S1**). To minimize misidentification errors, multiple comparisons of individual patterns were performed by a single observer using the pattern-matching programs Wild-ID (Bolger et al. 2012; http://www.dartmouth.edu/~envs/faculty/bolger.html) and Extract Compare (Hiby & Lovell 1990) in the three populations.

### MODEL 1: Modeling age-dependent survival in control, source, and relocated populations

We quantified age-dependent survival in control, source and relocated population using multievent CR models. In these models, a distinction is made between events and states (Pradel 2005). An event is the field observation coded in the individual’s capture history, which is related to the latent state of the individual. These observations can result in a certain degree of uncertainty regarding the individuals’ latent state. Multi-event models are designed to model this uncertainty in the observation process using hidden Markov chains (Pradel 2005).

We considered a model based on five states (**Supplementary material S2**) that include information about age (three age classes: juveniles ‘J’, subadult ‘S’, and adult ‘A’) and temporary emigration status at adult stage (available for capture ‘b’, and not-available ‘nb’). The models include four events coded in the capture histories as: ‘0’, individuals that are not captured; ‘1’, individuals captured as juveniles; ‘2’, individuals captured as subadults; and ‘3’, individuals captured as adults. Our model had robust design structure (Pollock 1982) so can consider temporary emigration. We considered both intra-annual and inter-annual emigration probabilities. Intra annual survival probability was set at 1 as is typical in robust design models (Kendall et al. 1995, Kendall & Nichols 2002). This assumption was realistic in our study system as intra-annual survival probability in *B. variegata* is usually > 0.9 (Cayuela et al. 2016a).

At their first capture individuals were assigned to one of three distinct states of departure (**Fig. 2**). At each time step, the information about individual state is progressively updated through four successive modeling steps: (1) survival, (2) age transition, and (3) temporary emigration. Each step is conditional on all previous steps. At the first modeling step, survival information is updated (**Fig. 3**). An individual may survive with a probability φ or die with a probability 1-φ. This results in a matrix with 5 states of departure and 5 states of arrival. Survival probability may differ between age classes by holding different φ values for the rows 1, 2 and 3-4. Note that the φ for the row 3 and 4 are always set equal as survival of adults that are available and unavailable for capture cannot be estimated separately due to parameter identifiability issues. In the second modeling step, information about individual age is updated (**Fig. 3**). An individual can change of age class with a probability α or remain in the same class with a probability 1-α. This leads to a transition matrix with 5 states of departure and 5 states of arrival. Note that in the matrix transition adults are forced to stay in the same age class. In the third modeling step, temporary emigration is updated. An adult may become available for capture with a probability γ or may become unavailable with a probability 1-γ. This results in a transition matrix with 5 states of departure and 5 states of arrival. The probability of capture availability at time *t* may depend on the individual status of capture availability at *t*-1 (i.e., Markovian emigration probability) by holding different γ values for the rows 3 and 4. The last component of the model (i.e., the event matrix, **Fig. 3**) links events to states. Individuals can be recaptured with a probability *p* or not with a probability 1-*p*. The probability of recapture may differ between age classes by holding different *p* values for the rows 1, 2, and 3. Note that the recapture probability of adults that are unavailable for capture is set at 0 in the matrix.

**Fig.3.**
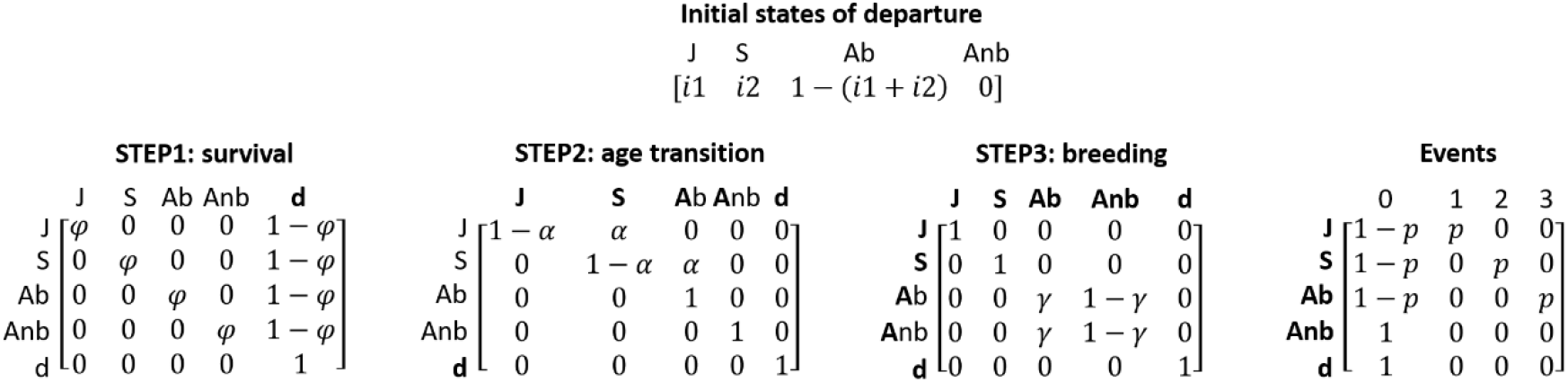
MODEL 1: state-state transition and events matrices. From a set of initial states of departure, individual state is updated through three successive modeling steps: (1) survival, (2) age transition, and (3) breeding attendance. The observation process is modeled through a single step (the matrix “Events”). The states and events are described in detail in Supplementary material 2, Table 1. In the state-state transition matrices, the departure states at time t-1 and the arrival states at time t are presented in rows and columns respectively. Whenever a status element is updated to its situation at t it appears in bold in the matrices (and stays bold throughout the following steps).

This parameterization was implemented in program E-SURGE (Choquet et al. 2009). The datasets from the three populations (control, source, and relocated) differed in terms of the number of study years and of capture sessions per year (**Supplementary material S1**). Therefore, we analysed the three datasets separately and considered one model selection procedure per population. We ranked models using the Akaike information criterion adjusted for a small sample size (AICc) and Akaike weights (*w*). If the Akaike weight *w* of the best supported model was less than 0.9, we used model-averaging to obtain parameter estimates (Burnham & Anderson 2002). The 95% CI were calculated using the delta-method (Royall 1986). Population-specific parameters were then compared based on their mean and their 95% CI. We examined our hypotheses about survival, temporary emigration, and recapture from the following general model: [φ(A), α(A), *γ_intra_*(S), *γ_inter_* (S), *p*(A + Y)]. The effects considered in the models were age (A), individual past emigration status (S) and year (Y). We hypothesized that: (1) survival φ probability differed among age classes (A), (2) emigration probability at both intra-annual (*γ_intra_*) and inter-annual (*γ_inter_*) levels depended on emigration status at *t*-1 (S), and (3) recapture probability varies according to age (A) and year (Y). We tested all the possible combinations of effects (including models with constant parameters noted ‘.’), resulting in the consideration of 16 competing models for the control and the relocated population (**Supplementary material S2**). For the source population, intra-annual and inter-annual emigration probability was set at 1 to avoid convergence issues, resulting in a restricted set (8) of competing models (**Supplementary material S2**).

### MODEL 2: Age-dependent survival of captive bred and locally born individuals

We aimed to compare survival of captive bred (and relocated individuals) and locally born individuals (i.e., the descendants of captive bred individuals) in the relocated population. In our study system, individuals released in the field cannot be identified with ventral coloration pattern. Therefore, there is an uncertainty about the ‘captive bred’ and ‘locally born’ status of individuals before they are juveniles (one overwintering). This status can only be ascertained for those that have been captured at juvenile and/or subadult stages (as we know their release or birth year in a given site). Because dispersal is rare between sites, it was possible to discriminate captive bred or locally born juveniles and subadults in each site. We used multievent CR models to take into account individuals’ status uncertainty in our survival inferences. We considered a model with nine states (**Supplementary material S2**) including information about age (juvenile, ‘J’; subadult, ‘S’; adult, ‘A’), the status ‘captive bred’ (‘T’) or ‘locally born’ (‘L’), and the emigration status (available for capture, ‘b’; non-available, ‘nb’). The model includes eight events that are coded as following in the capture histories: captive bred individuals are coded ‘1’ at juvenile stage, ‘2’ at subadult stage, and ‘3’ at adult stage; locally born individuals are coded ‘4’ at juvenile stage, ‘5’ at subadult stage, and ‘6’ at adult stage. We attributed the code ‘7’ to individuals whose status was uncertain and the code ‘0’ for individuals that were not captured.

As in the previous model, we considered three successive steps: (1) survival, (2) age transition, and (3) temporary emigration. In the first modeling, the information on survival is updated (**Fig. 4**). This results in a transition matrix with 9 states of departure and arrival. Survival probability φ may differ between the ‘captive bred’ and ‘locally born’ status by holding different φ values for the rows 1-4 and 5-8. Next, age transition is updated (**Fig. 4**), leading to a transition matrix with 9 states of departure and arrival. In the third modeling step, temporary emigration is modeled (**Fig. 4**), leading to a transition matrix with 9 states of departure and arrival. The probability of becoming available for capture γ may differ between captive bred and locally born adults by holding different γ values for the rows 3-4 and 7-8. For the events, we considered two distinct matrices (**Fig. 4**) describing the observation process conditional upon individuals’ underlying states. In the first matrix (9 states in row and 15 observations in columns), we introduced a set of intermediate observations describing the recapture process: captive bred and locally born individuals belonging to the three age classes can be recaptured with a probability *p* or not (‘ns’) with a probability 1-*p*. The probability of recapture may differ between captive bred and locally born individuals by holding different *p* values for the rows 1-3 and 5-7. In the second matrix (15 observations in row and 8 observations in column), individuals with an uncertain status are assigned to ‘captive bred’ and ‘locally born’ groups with a probability β and 1–β respectively.

**Fig.4.**
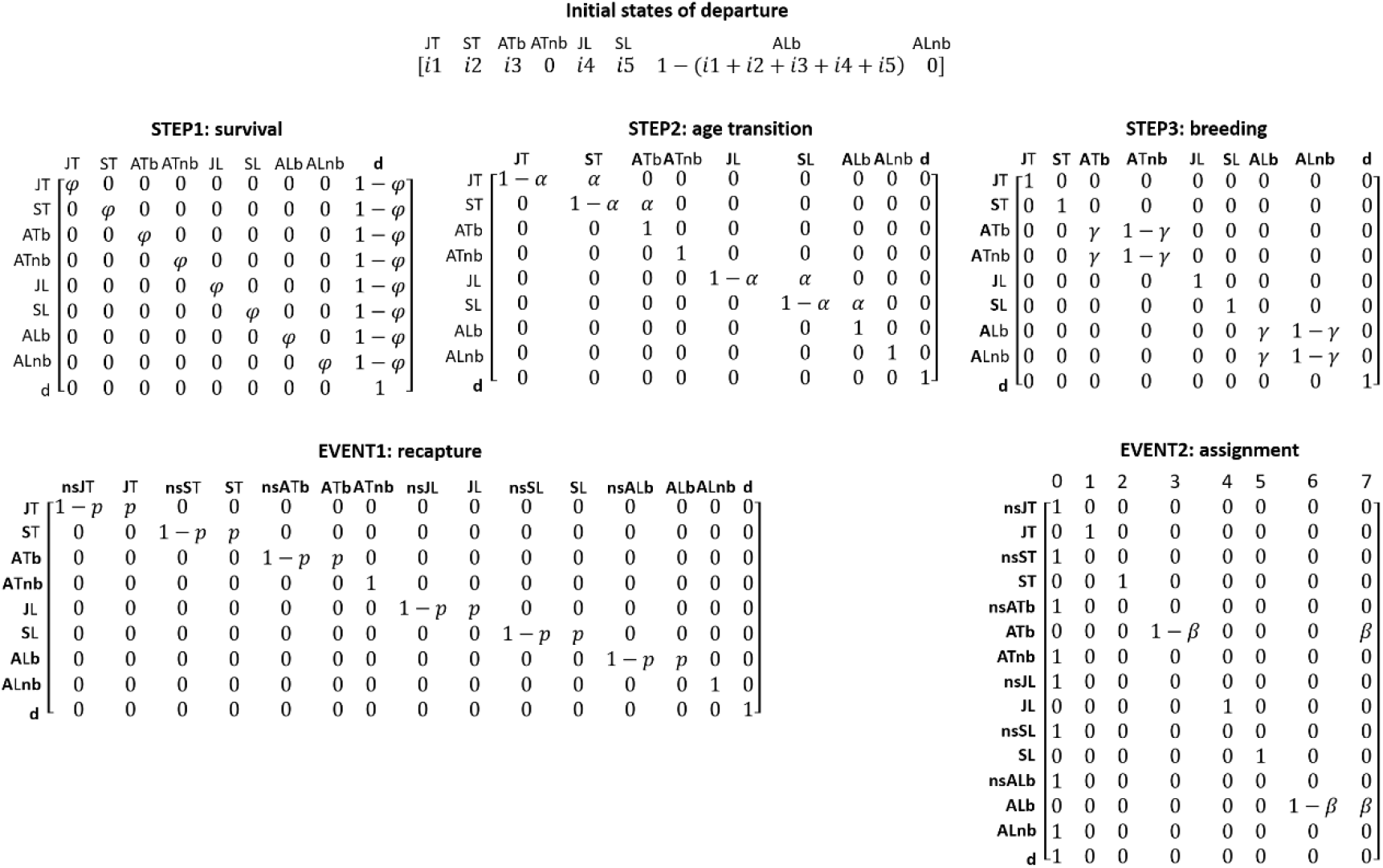
MODEL 2: state-state transition and events matrices. From a set of initial states of departure, individual state is updated through three successive modeling steps: (1) survival, (2) age transition, and (3) breeding attendance. The observation process is modeling through two successive steps: Event1 (recapture probability) and Event2 (assignment probability of individual with an uncertain status to ‘captive bred’ and ‘locally born’ groups. The states and events are described in detail in Supplementary material 2, Table 1.

This parameterization was implemented in program E-SURGE. The models were ranked using AlCc and AlCc weights (*w*). If the Akaike weight *w* of the best supported model was less than 0.9, we used model-averaging to obtain parameter estimates. We tested our hypotheses about survival, emigration probability and recapture from the following general model: [φ(A × T), α(A), *γ_intra_*(S × T), *γ_inter_*(S × T),*p*(A × T + Y), β(.)]. We considered three effects in the model: age (A), ‘captive bred’ and ‘locally born’ status (T), individual past emigration status (S), and year (Y). We hypothesized that (1) survival φ probability varied according to age class (A) and differed between captive bred and locally born individuals (T). As we were interested in obtaining the value of survival of both captive bred and locally born individuals (used in the simulation matrix model below), we kept the variable (T) in all the models. We considered an interaction between these two factors. We also examined whether emigration probability at both intra-annual (*Yintra)* and inter annual (*γ_inter_*) levels depended on emigration status at *t*-1 (S) and differed between captive bred and locally born adults; an interaction between these two factors was considered. We also hypothesized that recapture probability varied according to age (A), “captive bred” and “locally born” status (T), and year (Y); we considered an interaction between the factors ‘A’ and ‘T’ and added ‘Y’ in an additive way. Based on the results of the previous analysis (Model 1), we restricted the set of competing models. As the best-supported version of Model 1 was [φ(A), α(A), *γ_intra_*(S), *γ_inter_* (S), *p*(A + Y)] for the relocated populations, we kept this combination of effects as a baseline for Model 2. This led to the consideration of 8 competing models (**Supplementary material S2**).

### Modeling relocated population viability

We then examined the long-term viability of the relocated population by simulating population trajectories based on different scenarios. Based on the most realistic lifecycle for *B. variegata* (Cayuela 2015a, 2018b), we considered a three age-class (juvenile, subadult, adult), female-dominant, pre-breeding matrix (Caswell 2001) (**Fig. 5**). Fecundity *F* was possible for adults only and consisted of an estimation of recruitment, i.e., the number of recruited juvenile females at t per breeding female at *t*-1. We used the fecundity from the control population (*F* = 0.88; Cayuela et al. 2016) and another population from northeastern France (*F* = 0.79, Cayuela et al. 2018b). We used the age-dependent survival estimates resulting from Model 2 (*φ_juv_* for juveniles, *φ_sub_* for subadults, *φ_adu_* for adults). To be as realistic as possible, we considered environmental stochasticity for survival and fecundity rather than density dependence (Cayuela et al. 2018b). For each year, demographic parameter values were randomly sampled in a gaussian distribution centered on the estimates provided by the CR analysis and standard deviation inferred in two previous studies (Cayuela et al. 2016a, 2018b). The standard deviation values were: 0.05 for *φ_juv_*, 0.03 for *φ_sub_*, 0.01 for *φ_adu_*, 0.02 for *F*.

**Fig.5.**
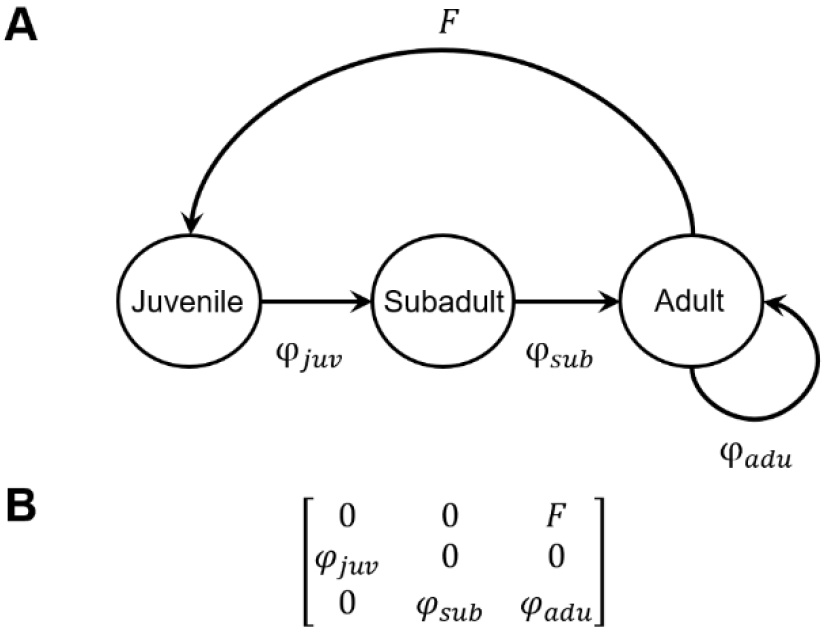
Lifecycle of Bombina variegata (A) and the simulation matrix model (B). The parameter φ juv corresponds to survival between the juvenile and subadult stages, φ_sub_ to survival between the subadult and adult stages, and φ_adu_ to survival at adult stage. The parameter *F* corresponds to female fecundity at adult stage.

Because earlier work indicates that relocated individuals can suffer loss of reproductive success (Saltz & Rubenstein 1995, Christie et al. 2014), in our simulations we considered a range of values for fecundity in the captive bred relocated individuals: 1) fecundity *F* is equal to 0.79 (control population) or 0.88 (the other population from northeastern France); and 2) a loss of 50% of the fecundity *F* is experienced. Each simulation began with 100 individuals. The number of individuals in each age class was obtained through the stable stage distribution provided by the three age-class Leslie matrix. The number of individuals at *t*+1 given the number of individuals at *t* was simulated using demographic stochasticity for captive bred individuals and their locally born descendants. We then simulated the number of individuals in 6 classes (relocated juveniles, subadults, adults; and locally born juveniles, subadults and adults). The number of surviving individuals in each age class and status was randomly sampled from a binomial distribution. The number of new recruits was sampled from a Poisson distribution. Offspring of captive bred individuals automatically became locally born individuals. The simulated population was monitored for 50 years and we performed 1000 simulations for each scenario. At each time step, we monitored the number of captive bred and locally born adults (i.e., breeding females).

## Results

### MODEL 1: Survival in control, source, and relocated populations

For the control and relocated populations, the best-supported model was [φ(A), α(A), *γ_intra_*(S), *γ_inter_* (S), *p*(A + Y)] (AICc weight = 1). For the source population, the best supported model was [φ(A), α(A), *γ_intra_* (0), *γnter* (0), *p*(A + Y)] (AICc weight = 0.83). Given the AICc value, we model averaged to obtain parameter estimates (**Supplementary material S2**). The recapture probability varied between populations, age, and year (**Supplementary material S2**). It was highest in the source population. The recapture probability was lower in subadults compared to juveniles and adults in all the populations.

Emigration probabilities in the relocated and the control population were Markovian. In both of these populations, at the intra-annual level, the probability of being available for capture at time *t* was higher for individuals that were already available at t-1 (control: 0.80, 95% CI 0.75-0.84; relocated: 0.70, CI 0.61-0.78) than for individuals that temporarily emigrated at *t*–1 (control: 0.06, 95% CI 0.03-0.12; relocated: 0.14, CI 0.10-0.18). At the inter-annual level, the pattern was the same: individuals that were already available for capture at *t*–1 had a higher probability of being available again at *t* (control: 0.64, 95% CI 0.59-0.69; relocated: 0.21, CI 0.11-0.34) than individuals that temporarily emigrated *t*–1 (control: 0.39, 95% CI 0.30-0.49; relocated: 0.13, CI 0.06-0.27).

Survival increased with age (**Fig. 6A**) in the source and control populations. There was a sharp increase from juvenile to subadult stages, then survival was relatively stable at subadult and adult stages. In contrast, in the relocated population, survival increased between juvenile and subadult stages and decreased at the adult stage. Importantly, our results showed that survival before sexual maturity (at juvenile and subadult stages) was relatively similar in the three populations and that adults experienced decreased survival in the relocated population.

**Fig.6.**
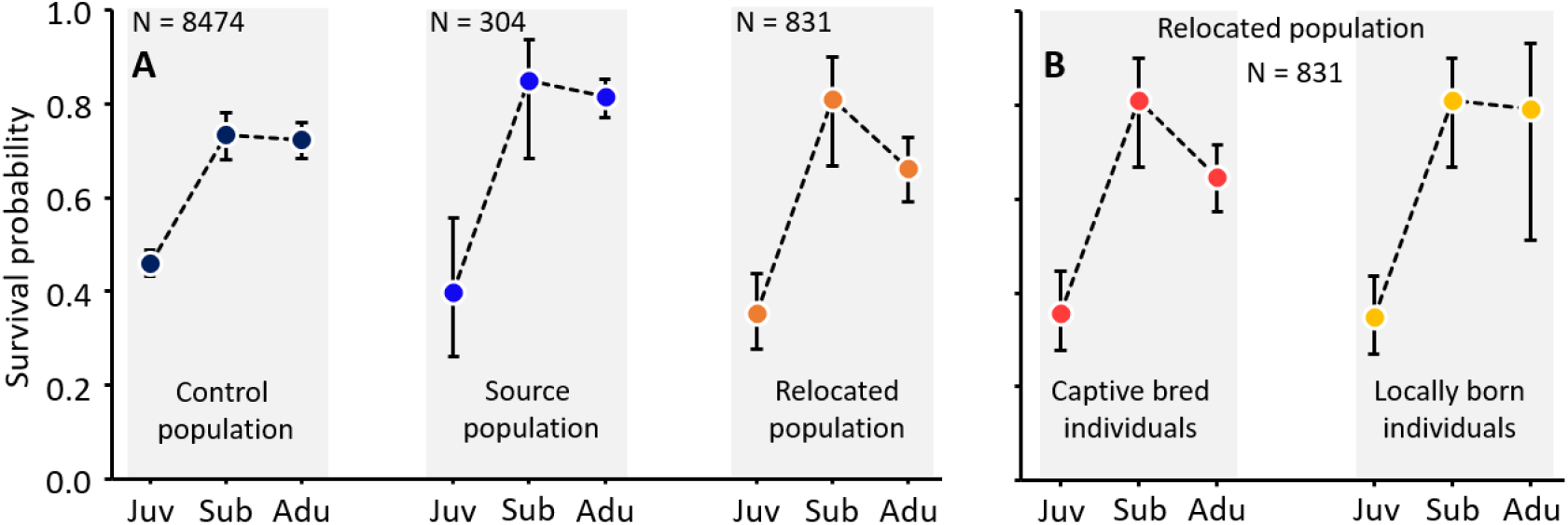
Age-dependent survival in control, source, and relocated populations of Bombina variegata. Three age classes are considered: juvenile, subadult, and adult. Before sexual maturity, survival pattern is relatively similar (i.e. overlapping 95% CI, showed as error bars) in the control, source and relocated populations (A). At adult stage, survival decrease in relocated population compared to the control and the source populations. In the relocated population, captive bred and locally born individuals also have a relatively similar survival regardless of individuals’ age (B).

### MODEL 2: Survival of captive bred and locally born individuals in the relocated population

The best-supported model was [φ(A×T), α(A), γ(S), p(A+Y), β(.)] (AICc weight = 0.51). Given the AICc value, we model-averaged to obtain parameter estimates. Overall, our results indicated that recapture probability marginally differed between captive bred and locally born-individuals and was similar to values provided by Model 1. As well, temporary emigration probability was similar in the captive bred and locally born individuals and equal to the values reported for MODEL 1 in this population. Survival of captive bred juveniles and subadults in the relocated population is similar to survival in their locally born descendants (**Fig. 6b**). In addition, survival of locally born adults was similar to survival of adults in the control and source populations (**Fig. 6a and 6b**). However, survival in captive bred adults was lower than survival in locally born adults (**Fig. 6b**). Although confidence intervals overlapped, the estimated means were quite different, 0.66 in captive bred adults and 0.79 in locally born adults.

### Simulation and long-term viability of the relocated population

Our simulations revealed that the relocated population is self-sustainable all the four scenarios considered (**Fig. 7**). In each of them, after 10 years captive bred individuals have disappeared from the population. The fecundity of captive bred and locally born individuals has a substantial effect on population trajectories: fecundity of 0.88 resulted in a substantial increase of the population size over time, while a fecundity of 0.79 led to a relatively stable population size over time. A reduction of the fecundity (−-50%) of captive bred individuals resulted in a reduced increase in the population size.

**Fig.7.**
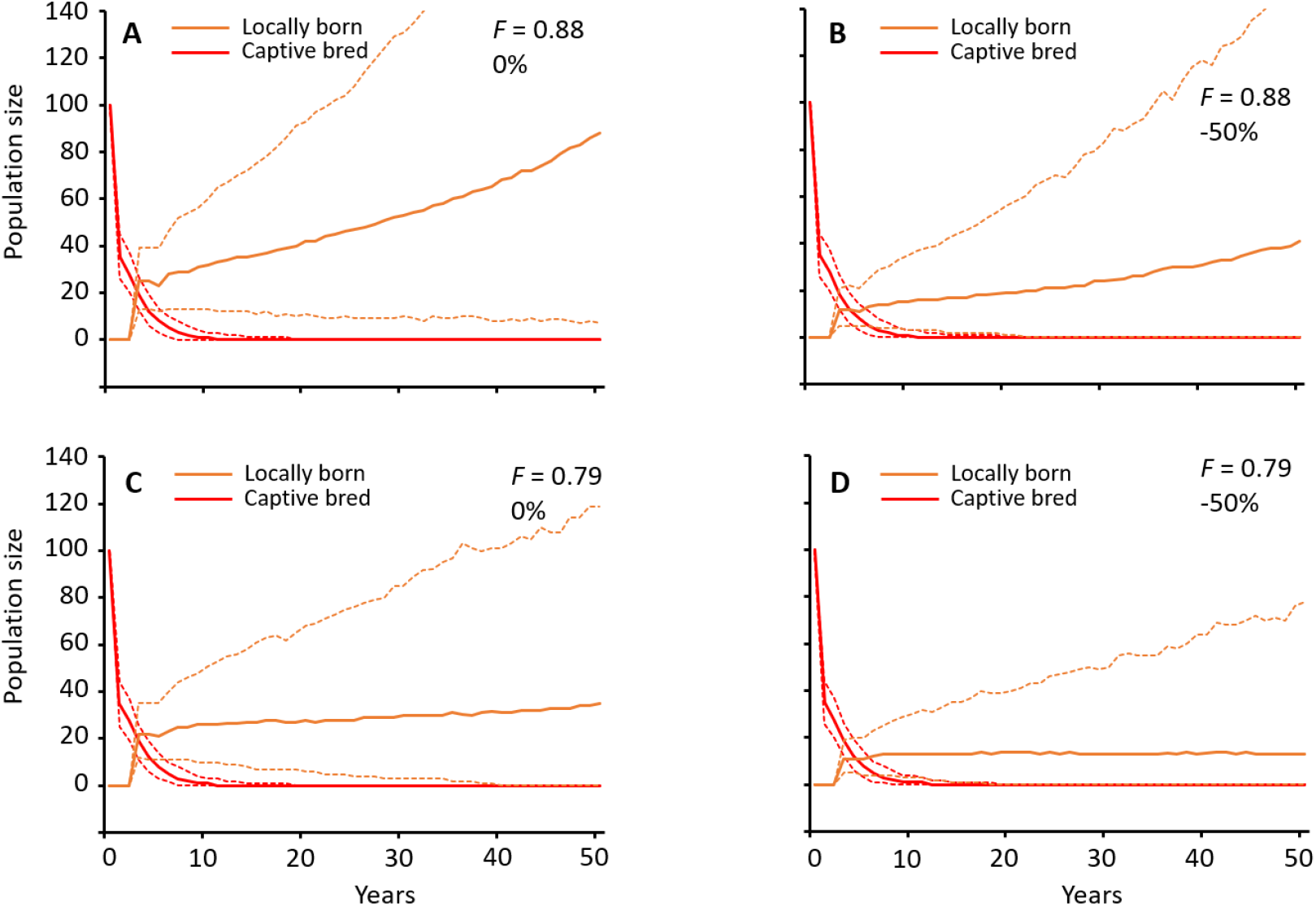
Simulated population size of captive bred and locally born individuals in relocated population of B. variegata over a 50-years period. We considered a range of situation where fecundity can be equal to that of population control (0.88; A) or that of another population from northeastern France (0.79; C). We examined the possibility that captive bred individuals may experience a fecundity decrease (−50%; B and C). Mean predictions are showed in full lines while their 95% CI are in dashed lines.

## Discussion

Our study showed that survival increased with age in the three populations. We also highlighted that the survival of adults from the relocated population was lower to that of individuals in the source and the control populations. This pattern resulted from a lower survival of captive bred individuals at the adult stage – the survival of their locally born descendants was similar that that reported in the control and source population. Furthermore, our simulations revealed that the relocated population should be self-sustainable in the four scenarios considered (i.e., no extinction). Yet, the fecundity drastically affected the fate of the population (i.e., stability or increase).

### Age-dependent survival pattern

We found that survival increases with age in *B. variegata* (except in adults that were captive bred) and that in general survival increases with age and becomes stable in adults. This pattern is congruent with data from other populations in Western Europe (Cayuela et al. 2016a, 2016b) and agrees with reports showing that post-metamorphic survival increases with body size in pond-breeding amphibians (Altwegg & Reyer 2003, Schmidt et al. 2012). Lower survival in smaller animals makes intuitive sense because small animals are likely more sensitive to dehydration and temperature variation, resulting in higher mortality (Hillman 2009). Our data are an important addition because they shows that this pattern remains consistent among distinct populations in different environmental contexts (forest, quarry) and with divergent demographic history (established and relocated).

### Post-relocation age-dependent mortality processes

Captive bred adult survival was lower in the relocated population than adult survival in source and control populations, but survival in captive bred juveniles and subadults was relatively similar in all three populations. Interestingly, by the time locally born descendants reached adulthood, there was no difference in survival rates compared to the other populations in any lifestage. In this case, relocation means a drastic change of environmental conditions for released individuals; the source population occupies a quarry dominated by a regrowth forest with artificial breeding pools, while the new habitat is made of ruts within a forest-grassland. The lack of performance of captive bred adults likely results from external mortality factors (competition, predation or physiological stress response to the novel habitat) rather than from their genetic background. Indeed, if maladaptation had led to a loss of survival-related performances at adult stage (e.g., thermoregulation, hydroregulation, osmoregulation, and locomotor capacity), lower performance should have been detected in their offspring after sexual maturity. Yet, in contrast to their parents born in captivity, locally born individuals do not experience a survival decrease at adult stage in the new forest-grassland habitat; we did not detect survival difference with individuals from the source population. In addition, the survival of the locally born adults (0.79) was close to that of individuals from forest (ranging from 0.71 to 0.75 in various populations of *B. variegata*; see Cayuela et al. 2016a) and grassland environments (0.77, Cayuela et al. 2016a), suggesting similar survival-related performances regardless of the environmental context. This suggests that, once the second generation is established, survival is similar to that in natural populations. As adaptation is unlikely to occur in only one generation with a reduced standing genetic variation (30 individuals involved in the breeding program), phenotypic plasticity likely allows individuals from quarry environments to accommodate new environmental conditions through acclimatization

Our results suggest that captive breeding and relocation may have delayed effects on individual performance (e.g., adult survival) rather than immediate effects following relocation (e.g., juvenile survival). The absence of a sharp decrease in survival during the establishment phase in this amphibian contrasts with the dramatic survival decrease reported in birds (Armstrong et al. 2017), mammals (Jule et al. 2008), reptiles (Roe et al. 2010, but see Bertolero et al. 2007) and fishes (Jonsson et al. 2003). This could be because of high mortality typically observed in *B. variegata* juveniles regardless of environmental context or provenance of individuals (Cayuela et al. 2016a). However, decreased survival after relocation may have occurred at the tadpole and metamorph stage (before the first over-wintering) when we were unable to identify individuals. Mortality is generally high in egg/tadpole and metamorph life stages in amphibians (Werner 1986, Vonesh & De la Cruz 2002) and mortality is usually the highest during the months following individuals’ release in other vertebrates (Hamilton et al. 2010, Armstrong et al. 2017). The number of juveniles recaptured (**Supplementary material S1**) relative to the number of tadpoles released in our study (**Fig. 2**) suggests significant mortality between these two stages. This pattern is congruent with data from relocated tadpoles and metamorphs of *Anaxyrus boreas* (weekly survival probabilities0.49 to 0.75, Muths et al. 2014). These circumstances preclude a general assessment of the effect of relocation on individuals’ survival before the first overwintering. Studies to compare survival of locally born and captive bred tadpoles and metamorphs are needed, especially because our simulations showed that the realized fecundity (including the survival at these two stages) of captive bred individuals could have a strong impact on the trajectories of relocated populations.

### Drivers of the relocation success and conservation advices

Our simulations showed that, despite decreased survival in captive bred adults, the relocated population should be self-sustainable (stable or increasing) in the four scenarios that we assessed. Simulations also revealed that variation in fecundity has a strong influence on the fate of the population. We showed that a fecundity value of 0.88 resulted in an increasing population size over time while a value 0.79 resulted in stable population size. Thus, potential loss of fecundity in captive bred individuals may slow down population growth. These results suggest that fecundity should be monitored during the establishment phase and that actions to mitigate decreases in fecundity should be developed.

We suggest the following measures to improve the chance of relocation success in pond-breeding amphibians:

1. *Relocate a large number of individuals.* We optimized the chance of relocation success by releasing a large number of individuals (7804 tadpoles, 18 subadults and adults over a 6-year stocking period). In their meta-analysis, Germano & Bishop (2009) showed that the success of amphibian relocation increases with the number of individuals and is maximal when it is higher than 1000 individuals.
2. *The ontogenic stage at which individuals are relocated matters.* We increased the chance of relocation success by releasing tadpoles at late stage of development (Gosner stages 42-45) and metamorphosed individuals. Muths et al. (2014) showed that age (and therefore size and mass) of tadpoles at the relocation time positively influence survival probability in *A. boreas*.
3. *The quality of the relocation area matters.* Our study suggests that, in the relocated population, second generation individuals do not experience decreased survival. Our simulations showed that relocation success was highly sensitive to variation in realized fecundity (i.e., fecundity and offspring survival until the juvenile stage). Fecundity could be increased by providing high-quality breeding sites (Barandun & Reyer 1997, Boualit et al. 2018, Cayuela 2018b). In this conservation project, we improved the chance of relocation success by selecting a protected relocation area, thereby avoiding threats contributing to decline in *B. variegata* (i.e., habitat loss and anthropogenic landscape fragmentation; Cayuela et al. 2015b, 2018b). In addition, human activities (i.e. military and forestry vehicle traffic) in the relocation area resulted in the creation and maintenance of suitable breeding waterbodies (i.e. ruts) for *B. variegata.*

### Conclusion

Our study is one of the few to provide a robust assessment of relocation success in an amphibian by examining survival at different ontogenic stages using individual data from multiple populations. Although the circumstances of the populations in this study are not typical of pond-breeding amphibians, conclusions are likely informative to more typical systems and are applicable to relocation efforts generally. Our results quantify age-dependent survival processes in the dynamics of a relocated population, a consideration that is relevant to the relocation success for most endangered vertebrates.

## Acknowledgments

The authors would like to thank P. Bily, M. Brialmont, B. Crombaghs, A. Deschaseaux, P. Dupriez, D. Gilson, P. Goffart, E. Graitson, J.-Y. Grenson, J. Huysecom, S. Liégeois, P. Lighezzolo, Y. Neus, H. Pirard, D. Rouvroy, S. Vanderlinden and H. de Wavrin for their active participation to the captive breeding, relocation and/or monitoring during this project. We also want to thank Fabien Lassus and Nina King-Gillies for their contribution to data collection in the control population.

